# Unraveling the plasmidome of *Helicobacter pylori*: an unexplored source of potential pathogenicity

**DOI:** 10.1101/2025.01.06.631533

**Authors:** Bradd Mendoza-Guido, Juan D. Romero-Carpio, Silvia Molina-Castro

## Abstract

2.

*Helicobacter pylori* is a significant human pathogen associated with gastric diseases, yet the role of plasmids in its pathogenicity remains underexplored. This study provides a comprehensive genomic and functional characterization of the *H. pylori* plasmidome using publicly available plasmid sequences. Of 335 plasmids analyzed, 127 non-redundant representatives were selected for downstream analysis. MobMess network and Mash distance-based analyses suggest two major plasmid groups (G1 and G2) with distinct genomic and functional profiles. G1 plasmids were enriched in genes related to the Fic protein family and ABC transporters, potentially contributing to pathogenicity through post-translational modification and antibiotic extrusion, respectively. In contrast, G2 plasmids were predominantly associated with the Type IV Secretion System (T4SS) and conjugative elements, indicating a role in horizontal gene transfer. Notably, compound genes were linked to pathogenic proteins, including ClpP, VapD, and a putative vacuolating cytotoxin-related gene. This study underscores the functional diversity of the *H. pylori* plasmidome and highlights the need for experimental validation to clarify its role in pathogenicity, antimicrobial resistance, and adaptability.

**Impact statement:** Although *Helicobacter pylori* is well-studied as the primary etiologic agent of gastric diseases, information about the role of plasmids in this species remains limited. This study contributes to microbial genomics by characterizing the genomic and functional diversity of the *H. pylori* plasmidome available in public databases. Our findings highlight the importance of monitoring and conducting experimental assays to better understand the role of plasmids in pathogenic bacteria like *H. pylori*, which may harbor genes related to pathogenicity and antimicrobial resistance.

**Data summary:** The authors confirm all supporting data, and protocols have been provided within the article or through supplementary data files. The code employed for bioinformatic analyses can be obtained from the corresponding author upon request.

## 5. Introduction

*Helicobacter pylori* is a Gram-negative bacteria prevalent in approximately 50% of the human population colonizing the gastric mucosa and causing chronic gastritis, which can progress to severe gastroduodenal pathologies [1, 2]. Though the last decades multiple elements associated with its pathogenesis have been identified, the most relevant being the presence of a 40-kb chromosomal region known as the Cag pathogenicity island (PAI) that encodes the effector protein CagA, insertable into the gastric epithelial cells through a type IV secretion system (T4SS) [3].

Considered naturally competent, *Helicobacter pylori* is one of the most genetically diverse pathogenic bacteria [4, 5]. Different studies have explored mobile genetic elements (MGEs) associated with this genus, focusing primarily on the analysis of the different T4SS present and DNA processing proteins [6–8]. Using publicly available databases, a chromosome painting and fineSTRUCTURE analysis clustered recombination events in *Helicobacter pylori*, providing insights into its population structure, extent and direction of genetic flux between subgroups [9]. Nonetheless, characterization and further analyses in the complete plasmidome of *H. pylori* is scarce.

Plasmids in bacteria represent a diverse and mobile source of adaptative and evolutionary traits. Understanding how the plasmidome influences fitness of pathogenic bacteria is crucial, as it contributes specific functional traits such as antibiotic resistance, virulence genes, biosynthesis of essential nutrients and modification of transfer RNAs [10]. This study aimed to characterize the sequences of all plasmids associated with *H. pylori* that were available in public databases, focusing on their genomic relationships and functional potential linked to pathogenic traits. Our findings revealed that *H. pylori* plasmids can be broadly categorized into two distinct groups with unique genomic and functional characteristics, both serving as potential reservoirs of pathogenic genes.

## 6. Methodology

### 6.1 Plasmid network and MASH comparison analyses

All plasmid sequences and metadata were downloaded from the PlasmidScope web server [11], selecting only plasmids with *Helicobacter pylori* as the host across all available databases (PLSDB, IMG-PR, COMPASS, GenBank, RefSeq, ENA, Kraken2, DDBJ, TPA and mMGE). Plasmids shorter than 1000 bp were excluded from the analysis. The remaining plasmid sequences were imported into Anvi’o v8.0 [12], where coding sequences were identified using Prodigal v2.6.3 [13] and annotated with COG v14 [14] and Pfam v32 [15]. Subsequently, we employed the PlasX-MobMess pipeline [10] to generate an evolutionary network of plasmid clusters and classify sequences into backbone and compound plasmids. The plasmid network was visualized with Cytoscape v3.10.3 [16].

MobMess defines a plasmid cluster as “compound” when it originates from a backbone cluster but contains additional “cargo” genes. Furthermore, the software classifies plasmids into fragment and maximal clusters. Fragment clusters are homologous to backbone clusters but lack circular plasmids, suggesting they may represent incomplete clusters that give rise to maximal clusters with “cargo” genes.

From each cluster, non-redundant plasmids (NR-plasmids) were selected through a dereplication step for further downstream analyses. In this process, all sequences presented into the same cluster are aligned in an all-versus-all manner using MUMmer [17], and the plasmid with the highest global sequence identity (Iglobal), averaged across all alignments, is chosen as the representative plasmid. In addition, only NR-plasmids belonging to plasmid networks with more than 2 clusters were selected (Figure 1).

**Figure 1.**
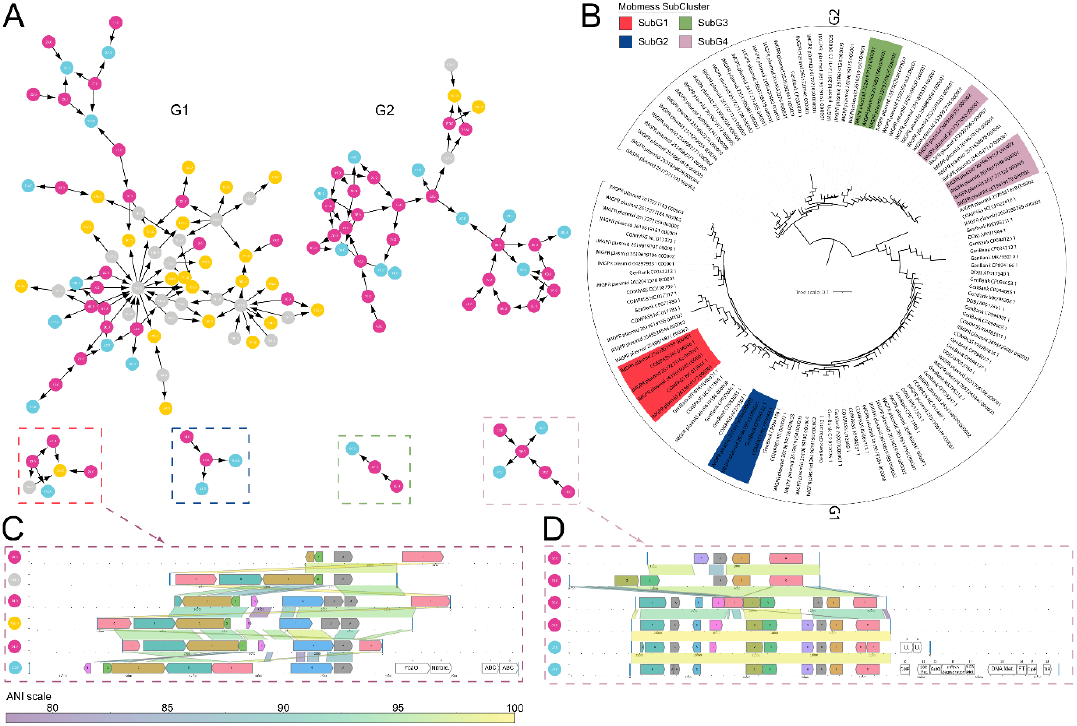
A) MobMess networks of *H. pylori* plasmid sequences. Nodes represent clusters containing redundant plasmids, connected by edges to other clusters with similar sequences. Node colors indicate plasmid categories: grey for backbone clusters, yellow for compound clusters (derived from backbone clusters), fuchsia for fragment clusters (backbone without any circular plasmid), and light blue for maximal clusters (derived from fragment clusters). The two main plasmid networks are labeled as G1 and G2, while additional networks are enclosed in red (SubG1), blue (SubG2), green (SubG3), and pink (SubG4) boxes. Numbers within nodes correspond to cluster IDs, with further details in Supplementary Table 4. B) Dendrogram based on MASH distances of NR-plasmid sequences, using a k-mer value of 21 and a sketch size of 10,000. Subgroups identified by MobMess are highlighted with the same colors as the boxes in A, while plasmids associated with G1 and G2 are marked with brackets. Alignments of plasmids in C) SubGroup-1 (SubG1) and SubGroup-4 D) (SubG4), ordered by gene content similarity (Jaccard index). Block colors represent shared family functions, and line colors correspond to ANI values shared between plasmid genes. Blank blocks represent genes unique to individual plasmids, with functions noted in text boxes: U = Unknown, OMP = Outer Membrane Protein, ABC = ABC transporter-associated protein, Nitrore = Nitroreductase, 23S = Adenine C2-methylase of 23S rRNA, GutQ = D-arabinose 5-phosphate isomerase GutQ, mRNA degradation = ribonuclease J1/J2, 16S Met = 16S dimethyltransferase RsmA/KsgA/DIM1, DNA Met = Adenine-specific DNA methylase, FT = Formyltetrahydrofolate hydrolase, ClpP = Periplasmic serine protease ClpP, and TIR = TIR domain. COG and Pfam annotations for all genes in SubG1 and SubG3 are available in Supplementary Files 2 and 3, respectively. High quality version of the figure is included at the end of the manuscript.

Plasmid sequence alignments of some NR-plasmids were visualized using the MobMess visualization function. Additionally, we constructed a dendrogram based on MASH distances for all NR-plasmid sequences to assess the genomic similarities among *H. pylori* plasmids using the mashtree software v1.4.6 [18] with a k-mer length value of 21 and sketch size of 10000.

### 6.2 Genomic characterization and functional annotation of non-redundant plasmids

Based on the MobMess network and MASH analyses, the NR-plasmids were divided into two groups, referred to as Group 1 (G1) and Group 2 (G2). We annotated replicase, relaxase and conjugative elements of NR-plasmid sequences (when available) with Mob-suite v3.1.8 [19]. Protein sequences from the NR-plasmids, obtained using Prodigal v2.6.3, were categorized into functional classes using the COGclassifier v1.0.5 software [20]. The distribution of coding sequences (CDS) across functional categories, along with GC content and sequence length for each group, were visualized using the ggplot2 library [21] in R software [22]. Additionally, a pangenome analysis of all NR-plasmids was conducted using Anvi’o v8.0, and a functional enrichment analysis [23] was performed to compare plasmids from G1 and G2, aiming to identify functions associated with gene clusters prevalent in each plasmid group.

Moreover, we classified genes as “backbone” if they were identified in plasmid sequences assigned to backbone or fragment clusters. Conversely, genes with COG accession numbers found exclusively in compound or maximal clusters (absent in backbone or fragment clusters), were categorized as “cargo” genes.

## 7. Results

### 7.1 Plasmid network and MASH comparison

A total of 335 *H. pylori* plasmid sequences obtained across 10 databases were retrieved, from where 12 were excluded due to lengths shorter than 1,000 bp. The remaining 322 plasmids were analyzed to construct a plasmid network using MobMess. Detailed information on these plasmids, including accession numbers, mobility classification, GC content, length, and topology, is provided in Supplementary Table 1. These plasmid sequences were grouped into 195 clusters, of which only 127 formed networks with more than two clusters and were retained. These 127 clusters were categorized as 22 backbone, 55 fragment, 25 compound, and 25 maximal clusters. Non-Redundant (NR) plasmids, selected as representatives for each cluster, were subsequently used for downstream analyses (Supplementary Table 2).

The plasmid network analysis revealed that most plasmid sequences are interrelated, forming two primary plasmid networks with clusters connected by edges, representing fragments shared among plasmid sequences (Figure 1A). Notably, most plasmid clusters in G2 are classified as fragment or maximal, except for two backbone clusters (177 and 131), which gave rise to compound clusters 180 and 181. This reflects the lack of circularity in the plasmid sequences of this group. Additionally, smaller plasmid networks were identified, isolated from the two main networks.

Likewise, the MASH distance-based dendrogram suggests that the plasmids forming the two main networks in the MobMess analysis (Figure 1B) can be categorized into two major genomic groups (Figure 1B). However, the small plasmid networks isolated in the MobMess analysis were found to fall within these two genomic groups in the mash dendrogram. Based on these findings, subsequent analyses focused on plasmid sequences categorized into Group 1 (G1) with 80 sequences and Group 2 with (G2) 47. Plasmids observed to be isolated from the main groups in the MobMess network were further designated as Subgroups 1 to 4 (SubG1–4) (Figures 1A and 1B). Of these four small networks, only SubG1 presented a cluster with circularity. The alignment of NR-plasmids of this network is visualized in Figure 1C.

Remarkably, the defined group classification is distinct from the Mash clustering analysis performed by the Mob-suite software, which uses a Mash distance threshold of 0.06 for cluster group assignment [24]. The cluster group assigned to each plasmid sequence by Mob-suite is provided in Supplementary Table 3 and may prove useful for future analyses and experiments aimed at exploring functional and genomic patterns within more specific groups.

### 7.2 Genomic characterization and functional annotation of non-redundant plasmids

From these 127 NR-plasmids, 41 were categorized as non-mobilizable due to the absence of relaxase genes (36 from G1 and 5 from G2), whereas 78 were classified as mobilizable (44 from G1 and 34 from G2). Notably, only eight plasmids contained conjugative elements, all of which belonged to G2. In general, the GC content of both groups was low, consistent with the GC content of *H. pylori*. G1 plasmids exhibited two GC content peaks around 34% and 36%, while G2 plasmids showed a more stable distribution centered around 32.5% (Figure 2A). Additionally, the length distribution per group revealed that G1 plasmids were predominantly shorter than 10,000 bp, with a few longer sequences. In contrast, G2 plasmids displayed a broader size range, spanning from 4,000 bp to 50,000 bp (Figure 2B).

**Figure 2.**
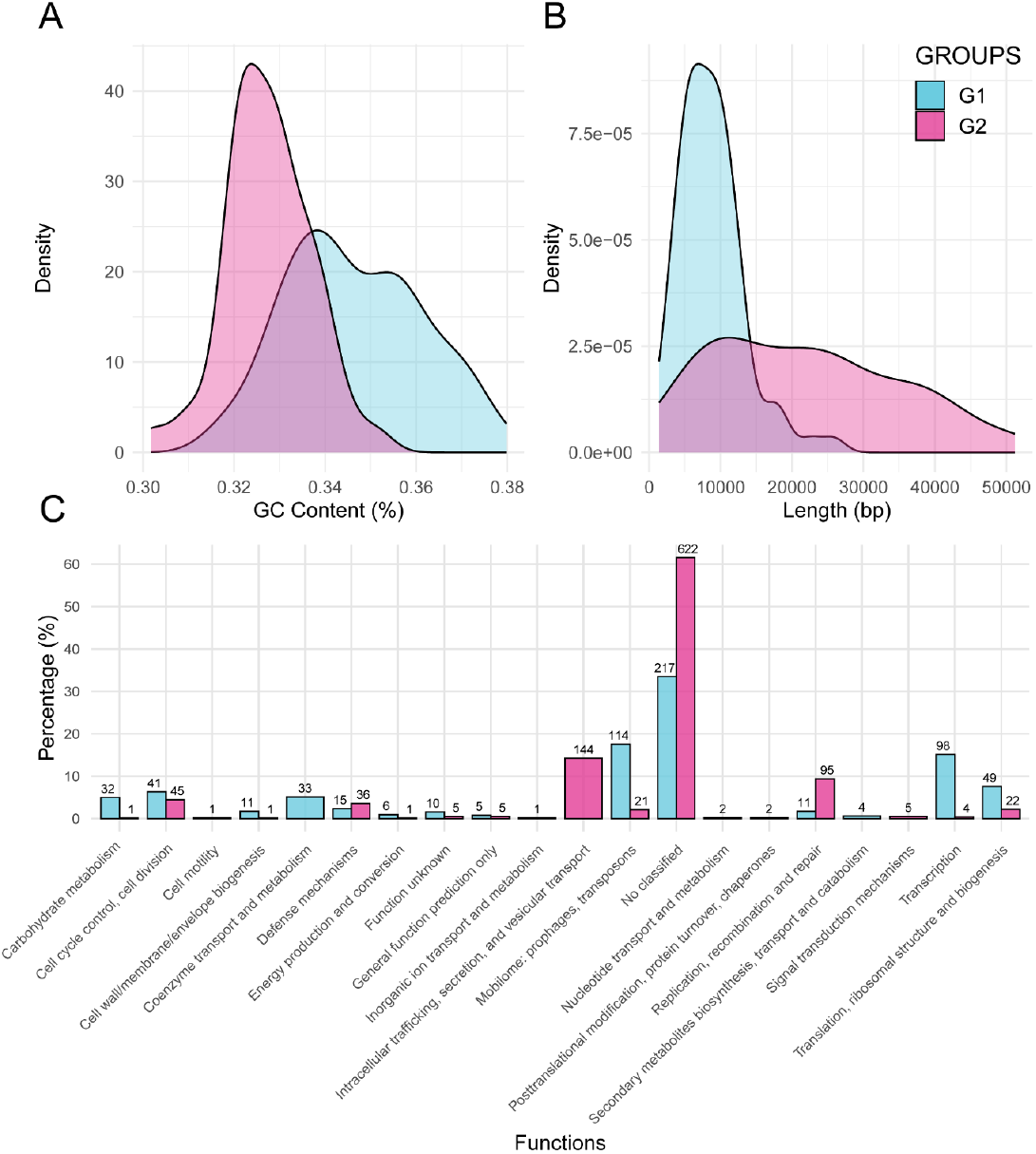
Distribution of A) GC percentage and B) plasmid length values observed in each proposed genomic group of *H. pylor*i plasmids. C) Percentage of Coding Sequences (CDS) assigned to different functional categories based on COG annotation for each plasmid group. The number above each bar indicates the number of CDS assigned to each COG category. Data for Group 1 is presented in light blue, and for Group 2 in fuchsia.

Regarding the functional traits observed in plasmid sequences, 217 CDS (33.49%) in G1 and 622 CDS (61.52%) in G2 could not be classified to any function. Plasmid sequences from G1 exhibited a higher proportion of CDS associated with transcription, translation, ribosomal structure and biogenesis, mobilome, and the metabolism of carbohydrates and coenzymes. In contrast, G2 plasmids were more strongly associated with intracellular trafficking, secretion and vesicular transport, as well as replication, recombination, and repair (Figure 2C).

A functional enrichment analysis of specific COG annotations revealed that G1 plasmids were primarily enriched for functions such as: Proteins involved in initiation of plasmid replication (mobilome) and Fic family proteins (transcription). Additional enriched functions in G1, albeit at lower prevalence, included Predicted arabinose efflux permease, MFS family (carbohydrate metabolism) and Molybdopterin or thiamine biosynthesis adenylyltransferase (coenzyme metabolism) (Table 1).

**Table 1.** COG functions enriched in each *H. pylori* plasmid group identified through functional enrichment analysis using Anvi’o v8.0. Only functions with an adjusted q-value below 0.01 are reported. The table includes the related function, enrichment score, adjusted q-value, and the prevalence of genes associated with each function in each plasmid group. Additionally, the analysis was performed using the Pfam annotation source, and the corresponding results are provided in Supplementary Table 5.

| COG14 function | Enrichment score | Prevalence in G2 (%) | Prevalence in G1 (%) |
| --- | --- | --- | --- |
| Type IV secretory pathway, VirD2 components (relaxase) | 118.56 | 98 | 1 |
| Protein involved in initiation of plasmid replication | 97.68 | 0 | 90 |
| ParA-like ATPase involved in chromosome/plasmid partitioning or Cellulose biosynthesis protein BcsQ | 92.30 | 81 | 0 |
| Superfamily II DNA or RNA helicase, SNF2 family or Adenine-specific DNA methylase or Adenine-specific DNA methylase, N12 class | 85.52 | 77 | 0 |
| Type IV secretory pathway, VirD4 component, TraG/TraD family ATPase | 69.81 | 66 | 0 |
| DNA topoisomerase IA | 63.96 | 62 | 0 |
| Fic family protein | 58.94 | 2 | 73 |
| Integrase | 45.30 | 47 | 0 |
| Type IV secretory pathway ATPase VirB11/Archaeellum biosynthesis ATPase | 42.83 | 45 | 0 |
| Type IV secretory pathway, VirB10 components | 35.70 | 38 | 0 |
| Type IV secretory pathway, VirB9 components | 33.41 | 36 | 0 |
| Predicted nucleotidyltransferase component of viral defense system | 28.95 | 32 | 0 |
| Type IV secretory pathway, VirB8 component | 28.95 | 32 | 0 |
| Type IV secretory pathway, VirB4 component | 28.95 | 32 | 0 |
| Predicted arabinose efflux permease, MFS family | 25.13 | 0 | 40 |
| Molybdopterin or thiamine biosynthesis adenylyltransferase | 20.15 | 0 | 34 |
| Serine/threonine protein kinase HipA, toxin component of the HipAB toxin-antitoxin module | 7.03 | 9 | 0 |
| REP element-mobilizing transposase RayT | 6.90 | 15 | 3 |

In contrast, G2 plasmids were enriched for genes encoding proteins associated with the Type IV secretion system (intracellular trafficking, secretion, and vesicular transport). Notably, the VirD2 protein was present in 97.87% of sequences in this group. Additional functions enriched in G2 included adenine-specific DNA methylase and DNA topoisomerase IA, both classified under replication, recombination, and repair category, as well as the ParA-like ATPase involved in chromosome/plasmid partitioning (cell cycle control and division). Each of these functions showed a prevalence exceeding 60% in G2 plasmid sequences (Table 1).

From all detected CDS, only 21 annotated functions were present in compound or maximal plasmids but absent in backbone and fragment plasmids, classifying them as “cargo” genes. The complete list of these genes, their functions, source of annotation and the corresponding plasmid sequences are provided in Table 2. Among these 21 cargo genes, eight were found in the plasmid IMGPR_plasmid_2609460337_000002 (G2), which includes functions related to nucleic acid methylation. This plasmid belongs to SubG4, and the complete alignment of this plasmid network is shown in Figure 1D (cluster 142), where cargo genes are represented by blank boxes. Additionally, other important functions related to pathogenicity, such as the “Virulence-associated protein VapD” and the “Putative vacuolating cytotoxin” were observed in plasmids from G1 and G2, respectively.

**Table 2.** Coding sequences classified as “cargo” genes were only present in plasmids belonging to compound or maximal clusters. CDS were annotated using two databases, COG14 and Pfam, to ensure better coverage of functional traits. The table presents the Gene ID, source, accession number, potential function, observed plasmids, and the corresponding plasmid group. A complete list of all identified CDS, along with their functions and amino acid sequences, is provided in Supplementary Table 4.

| Gene id | Source | Accession number | Functions | Plasmids | Group |
| --- | --- | --- | --- | --- | --- |
| 1004 | Pfam,<br>COG14 | PF13394.6, COG0820,<br>PF04055.21 | 4Fe-4S single cluster domain,<br>Adenine C2-methylase RlmN of<br>23S rRNA A2503 and tRNA A37,<br>Radical SAM superfamily | IMGPR_plasmid_2609460337_000002 | G2 |
| 1005 | Pfam,<br>COG14 | PF00571.28,<br>COG0517!!!COG0794,<br>PF01380.22,<br>PF13580.6 | CBS domain, CBS domain, D-arabinose 5-phosphate isomerase<br>GutQ, SIS domain | IMGPR_plasmid_2609460337_000002 | G2 |
| 1006 | Pfam,<br>COG14 | PF00753.27,<br>PF07521.12,<br>COG0595 | Metallo-beta-lactamase<br>superfamily, Zn-dependent<br>metallo-hydrolase RNA specificity<br>domain, mRNA degradation<br>ribonuclease J1/J2 | IMGPR_plasmid_2609460337_000002 | G2 |
| 1007 | COG14,<br>Pfam | COG0030,<br>PF00398.20 | 16S rRNA A1518 and A1519 N6-dimethyltransferase<br>RsmA/KsgA/DIM1 (may also have<br>DNA glycosylase/AP lyase<br>activity), Ribosomal RNA adenine<br>dimethylase | IMGPR_plasmid_2609460337_000002 | G2 |
| 1010, 844 | Pfam | PF00551.19 | Formyl transferase | IMGPR_plasmid_2609460337_000002,<br>IMGPR_plasmid_2551306581_000001 | G2 |
| 1011, 845 | Pfam | PF01343.18 | Peptidase family S49 | IMGPR_plasmid_2609460337_000002,<br>IMGPR_plasmid_2551306581_000001 | G2 |
| 106 | Pfam,<br>COG14 | PF01936.18,<br>COG1432 | NYN domain, Uncharacterized<br>conserved protein, LabA/DUF88<br>family | COMPASS_NC_017373_1 | G1 |
| 1220 | Pfam | PF07510.11 | Protein of unknown function<br>(DUF1524) | IMGPR_plasmid_2619619157_000002 | G2 |
| 1221 | Pfam,<br>COG14 | PF03235.14,<br>COG1479 | Protein of unknown function<br>DUF262, Uncharacterized<br>conserved protein, contains ParB-like and HNH nuclease domains | IMGPR_plasmid_2619619157_000002 | G2 |
| 1267 | Pfam | PF18304.1 | SabA N-terminal extracellular<br>adhesion domain | IMGPR_plasmid_2619619162_000001 | G2 |
| 153 | Pfam | PF09827.9 | CRISPR associated protein Cas2 | COMPASS_NC_019564_1 | G1 |
| 153, 1472 | COG14 | COG3309 | Virulence-associated protein<br>VapD (function unknown) | COMPASS_NC_019564_1,<br>IMGPR_plasmid_2619619196_000002 | G1 |
| 1596 | Pfam | PF03077.14 | Putative vacuolating cytotoxin | IMGPR_plasmid_2660238415_000001 | G2 |
| 230, 241, 392 | Pfam | PF05732.11 | Firmicute plasmid replication protein (RepL) | GenBank_CP032035_1, GenBank_MK795212_1 | G1 |
| 382 | Pfam | PF01719.17 | Plasmid replication protein | GenBank_MK795211_1 | G1 |
| 540 | COG14, Pfam | COG0732, PF01420.19 | Restriction endonuclease S subunit, Type I restriction modification DNA specificity domain | IMGPR_plasmid_2529293159_000001 | G2 |
| 563 | Pfam | PF02384.16 | N-6 DNA Methylase | IMGPR_plasmid_2529293163_000001 | G2 |
| 563, 792, 1103, 1563 | COG14 | COG0553, COG0827, COG4646, COG4983 | Superfamily II DNA or RNA helicase, SNF2 family, Adenine-specific DNA methylase, Adenine-specific DNA methylase, N12 class, Uncharacterized protein, contains Primase-polymerase (Primpol) domain | IMGPR_plasmid_2529293163_000001, IMGPR_plasmid_2548876501_000001, IMGPR_plasmid_2617271099_000001, IMGPR_plasmid_2660238415_000001 | G2 |
| 844, 1010 | COG14 | COG0788 | Formyltetrahydrofolate hydrolase | IMGPR_plasmid_2551306581_000001, IMGPR_plasmid_2609460337_000002 | G2 |
| 845, 1011 | COG14 | COG0616 | Periplasmic serine protease, ClpP class | IMGPR_plasmid_2551306581_000001, IMGPR_plasmid_2609460337_000002 | G2 |
| 849 | Pfam, COG14 | PF15919.5, COG1598 | HicB_like antitoxin of bacterial toxin-antitoxin system, Predicted nuclease of the RNase H fold, HicB family | IMGPR_plasmid_2551306581_000001 | G2 |

## 8. Discussion

In this study, we present a comprehensive characterization of the complete available *H. pylori* plasmidome, identifying two major plasmid groups with distinct genomic and functional traits. *H. pylori* is a significant human pathogen associated with a high disease burden worldwide, underscoring the critical importance of understanding the role of plasmids in its pathogenicity and adaptation to its host.

Plasmid sequences classified into the G1, were in general shorter than 10kb (Figure 2B) and with high prevalence of CDS related with plasmid replication protein, Filamentation Induced by cAMP (Fic) family protein amongst others. Proteins belonging to the Fic family have been reported to play an important role in the post-translational modification of bacterial and Eukaryotic proteins. Indeed, it has been demonstrated that some pathogenic bacteria can secrete Fic proteins as toxins to interfere in the function of host cells, as well as serve as components of toxin/antitoxin modules [25]. The presence of these family proteins in *H. pylori* has been reported in both chromosome [26] and plasmid sequences [27], but their potential role in its pathogenesis remains unexplored. The high prevalence of Fic proteins in G1 plasmids (and their absence in G2), along with their association with pathogenic traits and protein regulation, highlights the need for experimental studies to explore their role in *H. pylori* regulation and adaptive systems.

Other notable CDS presented in G1 plasmids are those with related functions to arabinose efflux permease (Table 1) and ABC transporters. The latter was not detected in enrichment analyses using COG, as the annotation software divided the CDSs into two different categories. However, using the Pfam database for annotation, ABC transporters were statistically significant, with a prevalence of 13% in G1 plasmids and 2% in G2 plasmids (Supplementary Table 5). Efflux pumps might play a crucial role in some multidrug resistance (MDR) bacteria by exporting antimicrobial molecules out of the cell. ABC transporters, well-characterized for their role in antibiotic resistance across various bacterial taxa, have been shown to play a major role in the development of MDR *H. pylori* strains [28]. In Figure 1C we evidence the presence of two CDS encoding ABC transporters in the maximal plasmid COMPASS_NC_019561_1 as part of the SubG1 plasmid network. This suggests that G1 plasmids may carry ABC transporter genes, potentially contributing to antibiotic resistance in *H. pylori*.

G2 plasmids exhibited larger sizes, which correlated with a higher number of coding sequences (CDSs); however, more than 60% of these CDSs were annotated with unknown functions (Figure 2C). This group was notably enriched with genes related to the Type IV Secretion System (T4SS), a well-established mechanism essential for the pathogenicity of *H. pylori*. Interestingly, *H. pylori* has been shown to harbor four chromosomally encoded T4SSs: Cag, ComB, Tfs3, and Tfs4 systems. While the Cag system has been the most extensively studied, particularly for its role in delivering the *cagA* oncoprotein, the presence of other systems is also significant due to their association with DNA mobilization[29]. The Tfs3 and Tfs4 systems are linked to conjugative elements, largely owing to the presence of *xer* recombinases and relaxases [29]. Supporting this, we identified the *xerD* recombinase gene in the G2 conjugative plasmids IMGPR_plasmid_2529292961_000001 and IMGPR_plasmid_2645727646_000001. This evidence suggests that these plasmids may represent integrating conjugative elements (ICEs) and warrant further investigation to elucidate the role of conjugative elements on the rise of genomic diversity in *H. pylori* strains.

Genes that were exclusively present in compound and maximal plasmids, classified as cargo genes, were also analyzed. These genes represent genomic and functional traits that may have been recently acquired within the plasmid sequences. The plasmid IMGPR_plasmid_2609460337_000002 from G2 contained 8 out of the 21 cargo genes observed in its structure. Notably, it included a gene related to the periplasmic serine protease ClpP, which belongs to a group of proteases playing an important role in the virulence of various bacterial pathogens [30]. Additionally, the plasmid contained a gene associated with RlmN, an enzyme involved in the methylation of the 23S ribosomal subunit. Experimental introduction of RlmN variants into *E. coli* has been shown to increase resistance to multiple antibiotics by inhibiting the endogenous C2 methylation of A2503 in 23S rRNA [31], noting the importance of monitor variants of these genes in human pathogens.

Another cargo gene linked to pathogenicity was the Virulence-associated protein D gene (VapD), which has been associated with virulence in multiple microorganisms, including *H. pylori* [32]. Although the role of the VapD protein in *H. pylori* pathogenicity is not fully understood, it has been proposed that it plays a crucial role in protecting *H. pylori* against the hostile conditions inside gastric cells during the colonization process [32].

Additionally, a gene related to a putative vacuolating cytotoxin was identified in the conjugative plasmid IMGPR_plasmid_2660238415_000001 from G2. A subsequent BLAST protein search revealed 100% identity with the vacuolating cytotoxin from *H. pylori*, covering 1,650 out of 2,941 amino acid residues (data not shown) suggesting the partial presence of the protein. The vacuolating cytotoxin protein, specifically the VacA protein in *H. pylori*, has been associated with an increased risk of peptic ulcers and gastric cancer [33], underscoring the importance of understanding the potential role of this and other cargo genes observed in plasmid sequences in the pathogenicity of *H. pylori*.

## 9. Conclusions

Taking together, our results provide a comprehensive genomic and functional characterization of the complete, non-redundant plasmidome of *Helicobacter pylori*. Our analysis identifies two main plasmid groups, each carrying key genes that may play crucial roles in the bacterium’s pathogenicity and adaptability. Furthermore, the detection of cargo genes associated with virulence factors such as ClpP, VapD, and vacuolating cytotoxins highlights the significant role of plasmids in the pathogenic potential of *H. pylori*. Additionally, the high prevalence of genes with unknown functions underscores the need for further experimental investigations to fully understand the functional impact of plasmid-encoded genes and their contribution to the adaptability and pathogenicity of *H. pylori* in humans.

## 11. Supplementary Materials

All supplementary tables are provided in Supplementary File 1.

Supplementary File 1. Excel file containing all supplementary tables mentioned in the manuscripts.

Supplementary File 2. Complete annotation of coding sequences (CDS) present in representative plasmids from SubG1, performed using the COG14 and Pfam databases.

Supplementary File 3. Complete annotation of coding sequences (CDS) present in representative plasmids from SubG3, performed using the COG14 and Pfam databases.

Supplementary Table 1. Information on all plasmids downloaded from the PlasmidScope web server with a length greater than 1000 bp.

Supplementary Table 2. List of representative plasmids selected by MobMess, including their cluster type (backbone, compound, fragment or maximal) and the assigned group according to MobMess and MASH analyses.

Supplementary Table 3. MOB-suite annotation results from representative plasmids, including details on plasmid typing, mobility, and associated replicon types.

Supplementary Table 4. Complete list of all coding sequences (CDS) identified by Prodigal, along with their annotations using the COG14 and Pfam databases.

Supplementary Table 5. Pfam functions enriched in each *H. pylori* plasmid group identified through functional enrichment analysis using Anvi’o v8.0. Only functions with an adjusted q-value below 0.01 are reported.

