## Supplementary Files 1-3 for "Unraveling the plasmidome of *Helicobacter pylori*: an unexplored source of potential pathogenicity": Supplementary File 2.pdf

| accession | accession_representative | contigs | COG14_FUNCTION | Pfam |
|---|---|---|---|---|
| A | COG3177 | 6 | COG3177:Fic_family_protein | PF02661.18:Fic/DOC_family |
| 0 | PF13575.6 | 6 | COG4403:Lantibiotic_modifying_enzyme | PF13575.6:Domain_of_unknown_function_(DUF4135) |
| 1 | PF03432.14 | 6 | ---- | PF03432.14:Relaxase/Mobilisation_nuclease_domain_ |
| 2 | PF05713.11 | 6 | ---- | PF05713.11:Bacterial_mobilisation_protein_(MobC) |
| 3 | COG1132 | 5 | COG1132:ABC-type_multidrug_transport_system,_ATPase_and_permease_component | PF00005.27:ABC_transporter PF13304.6:AAA_domain,_putative_AbiEii_toxin,_Type_IV_TA_system |
| 4 | COG5527 | 4 | COG5527:Protein_involved_in_initiation_of_plasmid_replication | PF01051.21:Initiator_Replication_protein |
| 5 | COG3041 | 4 | COG3041:mRNA-degrading_endonuclease_(mRNA_interferase)_YafQ,_toxin_component_of_the_YafQ-DinJ_toxin-antitoxin_module | PF05016.15:ParE_toxin_of_type_II_toxin-antitoxin_system,_parDE PF15738.5:Bacterial_toxin_of_type_II_toxin-antitoxin_system,_YafQ_ |
| 6 | PF01061.24 | 1 | ---- | PF01061.24:ABC-2_type_transporter |
| 7 | COG1131 | 1 | COG1131:ABC-type_multidrug_transport_system,_ATPase_component | PF13304.6:AAA_domain,_putative_AbiEii_toxin,_Type_IV_TA_system PF00005.27:ABC_transporter |
| 8 | COG0778 | 1 | COG0778:Nitroreductase | PF00881.24:Nitroreductase_family |
| 9 | COG1944 | 1 | COG1944:Ribosomal_protein_S12_methylthiotransferase_accessory_factor_YcaO | PF02624.16:YcaO_cyclodehydratase,_ATP-ad_Mg2+-binding |
