## Supplementary Files 1-3 for "Unraveling the plasmidome of *Helicobacter pylori*: an unexplored source of potential pathogenicity": Supplementary File 3.pdf

| accession | accession_representative | contigs | COG14_FUNCTION | Pfam |
|---|---|---|---|---|
| A | PF08843.11 | 6 | COG2253:Predicted_nucleotidyltransferase_component_of_viral_defense_system | PF08843.11:Nucleotidyl_transferase_AbiEii_toxin,_Type_IV_TA_system |
| 0 | COG3843 | 6 | COG3843:Type_IV_secretory_pathway,_VirD2_components_(relaxase) | PF03432.14:Relaxase/Mobilisation_nuclease_domain_ |
| 1 | COG0582 | 6 | COG0582:Integrase | PF00589.22:Phage_integrase_family PF02899.17:Phage_integrase,_N-terminal_SAM-like_domain |
| 2 | PF13155.6 | 5 | ---- | PF13155.6:Toprim-like |
| 3 | PF04610.14 | 5 | ---- | PF04610.14:TrbL/VirB6_plasmid_conjugal_transfer_protein |
| 4 | COG0550 | 4 | COG0550:DNA_topoisomerase_IA | PF01131.20:DNA_topoisomerase PF01751.22:Toprim_domain |
| 5 | COG1192 | 4 | COG1192:Cellulose_biosynthesis_protein_BcsQ | PF13614.6:AAA_domain PF01656.23:CobQ/CobB/MinD/ParA_nucleotide_binding_domain PF06564.12 PF09140.11:ATPase_MipZ PF10609.9:NUBPL_iron-transfer_P-loop_NTPase PF07015.11:VirC1_protein |
| 6 | COG3550 | 4 | COG3550:Serine/threonine_protein_kinase_HipA,_toxin_component_of_the_HipAB_toxin-antitoxin_module | PF07804.12:HipA-like_C-terminal_domain |
| 7 | PF08401.11 | 3 | ---- | PF08401.11:Domain_of_unknown_function_(DUF1738) |
| 8 | PF07510.11 | 1 | ---- | PF07510.11:Protein_of_unknown_function_(DUF1524) |
| 9 | COG1479 | 1 | COG1479:Uncharacterized_conserved_protein,_contains_ParB-like_and_HNH_nuclease_domains | PF03235.14:Protein_of_unknown_function_DUF262 |
| 10 | PF01856.17 | 1 | ---- | PF01856.17:Helicobacter_outer_membrane_protein |
| 11 | COG0820 | 1 | COG0820:Adenine_C2-methylase_RlmN_of_23S_rRNA_A2503_and_tRNA_A37 | PF13394.6:4Fe-4S_single_cluster_domain PF04055.21:Radical_SAM_superfamily |
| 12 | COG0517!!!COG0794 | 1 | COG0517!!!COG0794:CBS_domain!!!D-arabinose_5-phosphate_isomerase_GutQ | PF00571.28:CBS_domain PF01380.22:SIS_domain PF13580.6 |
| 13 | COG0595 | 1 | COG0595:mRNA_degradation_ribonuclease_J1/J2 | PF00753.27:Metallo-beta-lactamase_superfamily PF07521.12:Zn-dependent_metallo-hydrolase_RNA_specificity_domain |
| 14 | COG0030 | 1 | COG0030:16S_rRNA_A1518_and_A1519_N6-dimethyltransferase_RsmA/KsgA/DIM1_(may_also_have_DNA_glycosylase/AP_lyase_activity) | PF00398.20:Ribosomal_RNA_adenine_dimethylase |
| 15 | COG0827 | 1 | COG0827:Adenine-specific_DNA_methylase | ---- |
| 16 | COG0788 | 1 | COG0788:Formyltetrahydrofolate_hydrolase | PF00551.19:Formyl_transferase |
| 17 | COG0616 | 1 | COG0616:Periplasmic_serine_protease,_ClpP_class | PF01343.18:Peptidase_family_S49 |
| 18 | PF13676.6 | 1 | ---- | PF13676.6:TIR_domain |
